# Ancient sedimentary DNA shows more than 5000 years of continuous beaver occupancy in Grand Teton National Park

**DOI:** 10.1101/2024.12.05.627036

**Authors:** D. Nevé Baker, Darren J. Larsen, Emily Fairfax, Amelia P. Muscott, Beth Shapiro, Sarah E. Crump

**Author notes:** Corresponding author: D. Nevé Baker.

## Abstract

Beaver-based restoration is emerging as a cost-effective conservation and climate adaptation strategy, but efforts are constrained by limited knowledge of pre-colonial beaver distribution and their long-term ecosystem impacts. Here, we apply sedimentary ancient DNA (sedaDNA) techniques to investigate the history of beaver occupancy at three lakes in Grand Teton National Park, Wyoming over the last ∼10 ka, as well as interactions with the local plant community. We documented a dynamic history of beaver presence in two sub-alpine lakes (Taggart and Jenny Lakes) and demonstrate no history of beaver occupancy at the higher-elevation alpine lake (Lake Solitude). Beavers were first detected at Jenny Lake around 7.2 ka and intermittently thereafter. At nearby Taggart Lake, beavers were first detected at ∼5.9 ka and continuously from 5.2 ka onwards. Vegetation metabarcoding revealed a shift in plant community coinciding with beaver establishment in these two sub-alpine lakes, as well as an increase in taxonomic diversity. These changes coincide with regional trends towards wetter conditions. Notably, beavers persist at Taggart Lake during inferred droughts, indicating a potential role in maintaining wetlands through extended periods of climatic stress. Our results demonstrate sedaDNA as a powerful, novel technique for reconstructing past beaver occupancy dynamics.

## 1. Introduction

As climate change intensifies, so does our need to find low-cost, sustainable ecosystem conservation and restoration solutions. Beaver-based restoration, which entails encouraging beaver establishment in low-functioning watersheds through both reintroductions and beaver mimicry, is one solution that is rapidly gaining momentum. Beavers are ecosystem engineers; by damming streams, beavers retain water on the landscape and slow sediment transfer, significantly altering the hydrology, geomorphology, and ecological community of riparian systems (Naiman et al. 1988, Gurnell 1998, Hood and Bayley 2008, Fairfax and Small 2018, Puttock et al. 2021). Beaver-modified landscapes support greater habitat complexity and diversity than their non-beaver occupied counterparts, and are more resilient to flood, drought and wildfire (Rosell et al. 2005, Mitchell and Cunjak 2007, Stringer and Gaywood 2016, Fairfax and Small 2018, Fairfax and Whittle 2020). However, questions remain as to where beaver reintroduction is appropriate, how beaver engineering affects the local environment at long (beyond decadal) timescales, and where beavers can survive and thrive in the future as land use patterns and local climates continue to change. These questions arise in part from a lack of understanding of the distribution and extent of historic beaver activity. Extensive trapping in the 17th-19th centuries depleted beaver populations throughout much of North America, bringing the species close to extinction (Wohl 2001). Most evidence of the range and distribution of beavers prior to fur trade decline is sociocultural, based largely on Traditional Ecological Knowledge, historical records from fur trappers, and indirect information such as place names (Lanman et al. 2012, 2013, Tape et al. 2021, Richmond et al. 2021). Physical evidence of beavers such as fossils, woody debris, and sedimentary proxies for dam building tends to be sparse and stochastically distributed (Robinson et al. 2007, Persico and Meyer 2009, 2013, Kramer et al. 2012, Mitchell et al. 2016, Davies et al. 2022). The lack of empirical data hinders efforts to reintroduce beavers to historically occupied regions and also limits scientific understanding of how beaver activity influenced long-term ecological and geological processes prior to European colonization of North America (Kramer et al. 2012).

Analysis of environmental DNA isolated from ancient sediments (sedaDNA) is a relatively new type of physical evidence used to understand past ecosystems, facilitated by recent advances in ancient and degraded DNA methodologies (Rawlence et al. 2014, Capo et al. 2021, Crump 2021). sedaDNA can be used to determine presence of specific taxa as well as to characterize ecological communities; alongside other paleosedimentary analyses, sedaDNA provides a powerful tool for reconstructing past environments and understanding the interaction between geological and ecological processes over long timescales (Thomsen and Willerslev 2015, Graham et al. 2016, Parducci et al. 2017, Crump 2021).

Here, we analyze sedaDNA from three lake sites in the Teton Range, Wyoming to document the historical presence of beavers in Grand Teton National Park (GTNP) over the last 10 ka and investigate beavers’ potential impact on local vegetation communities. GTNP and the greater region - including Yellowstone National Park - is one of the few areas in North America where Holocene beaver activity has been reconstructed from sedimentary analyses, making it an ideal location for testing this novel method. Sedimentary proxies from multiple stream beds in this region indicate sporadic beaver activity in the early Holocene and fairly consistent activity in the later Holocene with notable gaps corresponding to periods of regional drought and climatic anomalies (Persico and Meyer 2009, 2013). Beavers in the greater GTNP region were heavily trapped in the early 1800s but rebounded in the 20th century. As of 2014, 83 active beaver lodges were documented in GTNP, representing an estimated 400 individuals (Collins 1976, Gribb and Harlow 2014). The purpose of our study is twofold: 1) to demonstrate the utility of sedaDNA for documenting past presence of beavers in mountain watersheds, and 2) to enrich the current understanding of Holocene ecological dynamics in GTNP by reconstructing the local history of an environmental engineer and its interaction with plant diversity and community structure on geological time scales.

## 2. Methods

### 2.1. Study area

The Teton range, northwest Wyoming, is a fault-block mountain range located at the headwaters of the Snake-Columbia river network and within the Greater Yellowstone Ecosystem (Love et al. 2003). Its iconic topography forms the centerpiece of GTNP, which attracts over 3 million visitors annually, and is attributed to uplift along the Teton normal fault (Zellman et al., 2019) and periodic glaciations (e.g. Pierce et al., 2018). The tectonic and glacier-geomorphic history of the mountains created a series of adjacent glacial valleys carved into the steep eastern range front (Fig. 1). Each valley spans >1,500 vertical meters of elevation and transects multiple vegetation environments from high alpine tundra down to mixed conifer forests at the valley floor. Many of the valleys contain small alpine lake basins, positioned in high-elevation headwater cirques, which drain into a larger, moraine-dammed piedmont lake at the valley mouth (Fig. 1). Teton lakes preserve a continuous and datable sedimentary record of environmental conditions in the region since deglaciation approximately 15,000 years ago (e.g. Larsen et al., 2016).

**Figure 1:**
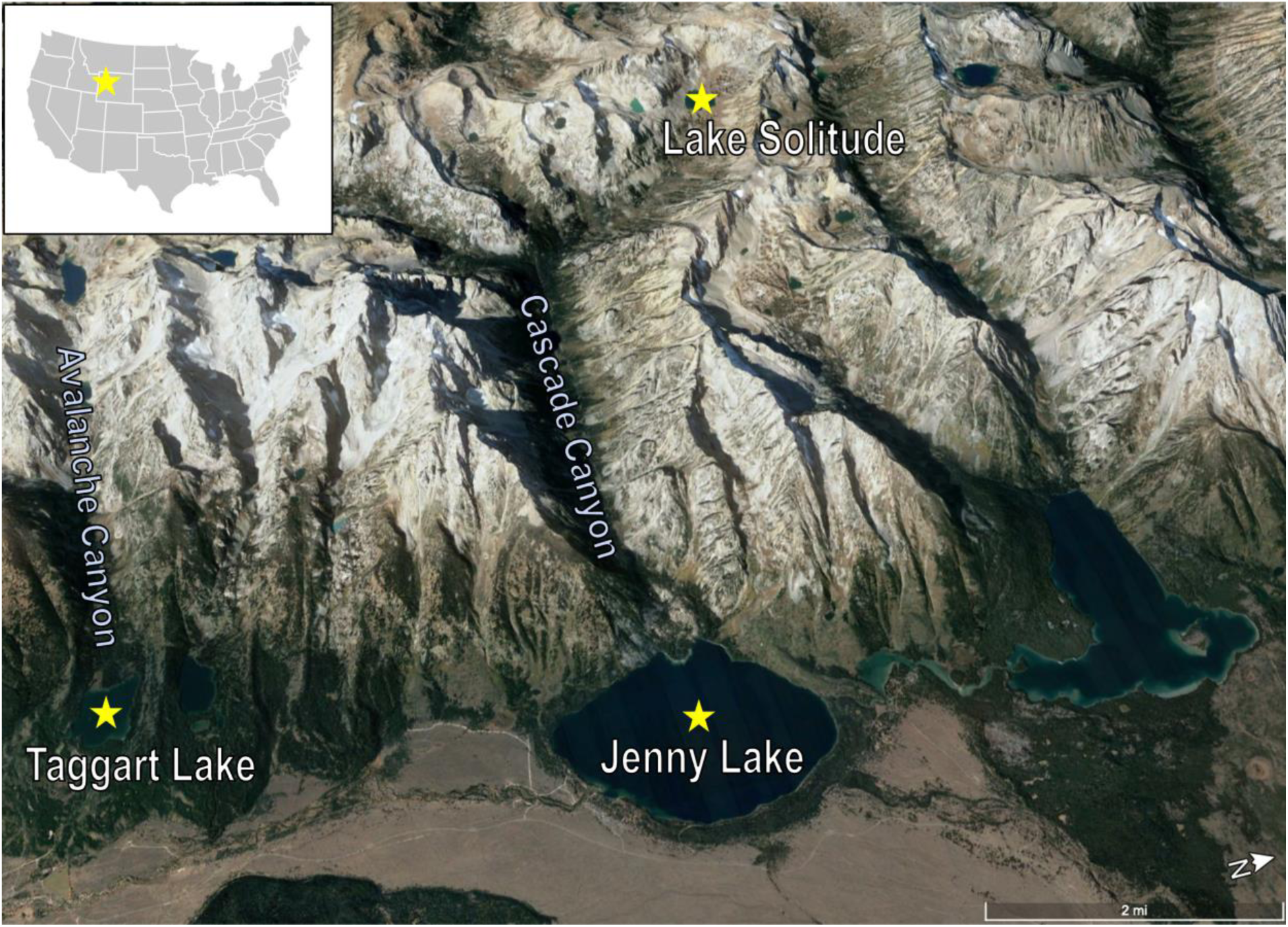
Oblique aerial view of the central Teton Range, Grand Teton National Park, showing the locations of the three lake sites (starred) included in this study. Inset map shows location of GTNP.

To determine the past presence of beavers at GTNP, we analyzed the sediment fill contained in three lakes in the central Tetons — Taggart Lake, Jenny Lake, and Lake Solitude (Fig. 1; Table S1). These lakes were chosen for their varying characteristics (size, depth, elevation) and beaver habitat suitability. Additionally, they are positioned in a region with a well-described geomorphic and paleoclimatic history (e.g. Whitlock, 1993; Larsen et al., 2016, 2020). Taggart and Jenny Lakes both occupy terminal basins excavated by valley glaciers during the last glaciation and are located at similar elevations at the mouths of their respective valleys. Jenny Lake (surface area: 5 km^2^, max depth: 73 m, elevation: 2070 m asl) is larger, deeper, and colder than Taggart Lake (surface area: 0.5 km^2^, max depth: 9.5 m, elevation: 2105 m asl) and has a larger catchment, suggesting a different aquatic profile. The moraine ridges and hillslopes surrounding both lakes support primarily mixed conifer forests. The creeks draining into Taggart and Jenny Lakes contain good beaver habitat and contemporary beaver activity in this area is well documented (Gribb and Harlow 2014, GBIF Secretariat 2022). Furthermore, sedimentary evidence indicates periodic beaver activity in this region of the Teton range front throughout the Holocene (Persico and Meyer 2013). Lake Solitude (surface area: 0.15 km^2^, max depth: 10.5 m, elevation: 2760 m asl) is located approximately ∼700 m higher than Taggart and Jenny Lakes, near the elevation of modern tree line and the top of the watershed that drains into Jenny Lake (Fig. 1). The lake occupies a small bedrock depression in a glacial cirque at the head of the north fork of Cascade Canyon. (Fig. 1). High snowfall and long, cold winters at this site result in prolonged seasonal lake ice cover for up to 9 months per year. At present, Lake Solitude is considered marginal beaver habitat and has no known history of beaver activity.

### 2.2. Sediment collection and age control

Sediment cores were collected from the central basins of each lake using multiple types of piston corers deployed from the lake surfaces (see Supplementary Information for details of field and analytical methodology; Table S1). Following collection, all cores were split, imaged and archived using conventional methods (see Supplementary Information). For each lake, we created a composite stratigraphic sequence and established a secure geochronology using radiocarbon dating of terrestrial plant macrofossils (e.g., conifer needles, charcoal, and woody plant fragments) and the position of the Mazama ash layer (∼7.6 ka) (Table S2; Egan et al., 2015, Larsen et al. 2016). The radiocarbon results were calibrated and converted to calendar years before present using CALIB 8.2 with the IntCal20 calibration curve (Stuiver et al. 2010, Reimer et al. 2013). Age-depth models were constructed for all lake sequences through interpolation of individual control points using the ‘classical’ age modeling code for R software (Figure S1; Blaauw 2010).

### 2.3. sedaDNA extraction and analysis

Core subsampling, extraction, and laboratory analysis was performed in dedicated ancient DNA clean rooms following standard ancient DNA protocols including full personal protective equipment and extensive bleaching of surfaces and tools. Two replicate 500mg subsamples were taken from the interior of the archived core half to minimize contamination and digested in a buffer following Grealy et al. (2015). One negative extraction control was prepared for each batch of 11 samples and included in all downstream analyses. Sediment digests were concentrated in Vivaspin centrifugal concentrators (Sigma Aldrich), added to a binding buffer following Dabney et al. (2013), and purified via MinElute PCR Purification Kit (QIAGEN). To evaluate inhibition and inform downstream analyses, extracts were first amplified via quantitative PCR (qPCR) with *trn*L barcode primers (Table 1) and a serial dilution (full concentration, 1/10, 1/100). qPCR results were used to inform sample-specific dilutions and target amplification cycles (cycle number at which exponential amplification ended) for the metabarcode library PCR.

**Table 1:**
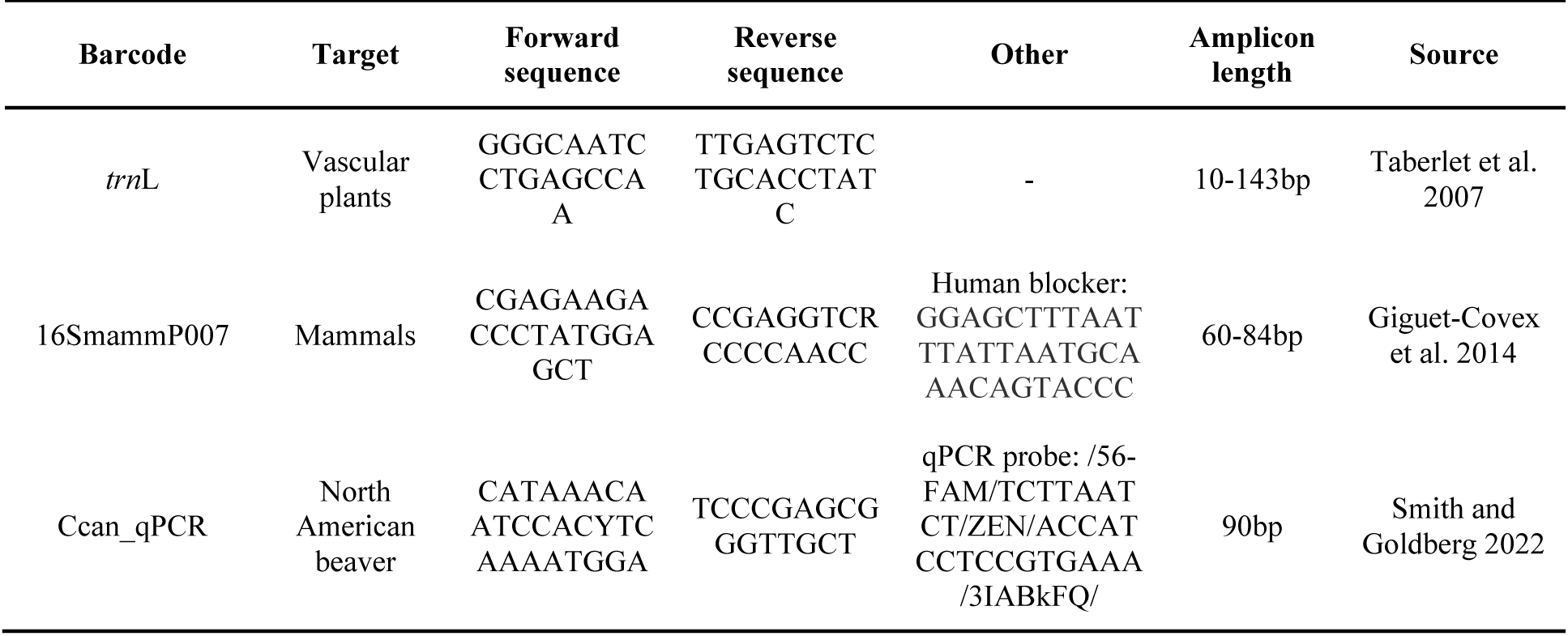
Primers used for qPCR and metabarcoding.

North American beaver (*Castor canadensis*) presence was assessed with targeted sequence detection through qPCR using a species-specific primer-probe assay developed by Smith and Goldberg (2022) that amplifies a 90bp fragment of the beaver mitochondrial genome (Table 1). We performed five replicate qPCRs for each extract (including controls) at the recommended dilution from the metabarcoding qPCR. Positive beaver detection was indicated by exponential amplification over a baseline threshold of 1000 relative fluorescence units (RFUs).

To investigate change in the vascular plant community, extracts were PCR amplified using barcode primers targeting the *trn*L P6 loop of the plant chloroplast genome with five replicates for each extract (Table 1). A barcode targeting the 16S region of the mammalian mitochondrial genome (16SmammP007) was also amplified and sequenced from all Jenny Lake and Taggart Lake samples to validate the beaver presence results from the species-species qPCR assay. Metabarcode libraries were generated using a two-step protocol (Nichols et al. 2018) with an initial metabarcoding PCR followed by a second indexing PCR to attach unique dual indexing primers. Libraries were quantified via Qubit Fluorometric Quantitation (ThermoFisher Scientific) and QIAxcel fragment analyzer (QIAGEN) to determine concentration and average fragment length and pooled in equimolar volumes for sequencing on an Illumina NextSeq 2×150 run, aiming for 50,000 reads per library. Sequencing reads were trimmed and processed with the Anacapa QC pipeline, then clustered as Amplicon Sequence Variants (ASVs) and ASVs assigned to taxa with the Anacapa CRUX pipeline (Curd et al. 2018). ASV assignments with a 60% Bayesian Confidence Cutoff were retained, following established methods (Curd et al. 2018, Lin et al. 2021). We used the decontam package in R (v1.12) (Davis et al. 2018) to compare taxonomic composition of samples and negative controls and remove any observed contamination. Following filtering, replicate libraries were merged, samples with fewer than ten reads were removed, and raw ASV counts for each sample were converted to relative abundance for downstream analyses of taxonomic abundance and beta diversity. Taxonomic abundance was visualized with the Phyloseq R package (McMurdie and Holmes 2013). Alpha diversity analyses were performed on unmerged sample replicates with Phyloseq. Compositional change in taxonomic assemblages was assessed using nonmetric multidimensional scaling (NMDS) ordination based on Jaccard similarity of the relative abundance data using the vegan R package (Oksanen et al. 2019).

## 3. Results

### 3.1. Sediment chronologies and sedaDNA

Teton lakes formed following regional deglaciation and their sediment fill spans the past ∼15 ka (Larsen et al., 2016). In this study, we focus our analyses on the past ∼10 ka common to all three lake core sequences. During this time interval, the average time between age-control points at Taggart Lake, Jenny Lake, and Lake Solitude is approximately 1,100, 900, and 1,000 years, respectively (Table S3; Fig. S1). Using the individual age models as a guide, we extracted and analyzed a total of 51 sedaDNA samples from the three lake cores; 20 each from Taggart Lake and Jenny Lake, and 11 from Lake Solitude (Table S3). Vascular plant sequencing using the *trn*L barcode yielded an average 259,541 reads and 50 identified genera per sample after quality filtering and merging replicates. 16SmammP007 libraries were generated for 9 samples from Taggart Lake and 15 from Jenny Lake, with an average of 36,732 reads and 4.5 identified genera per sample.

### 3.2. Beaver detection

The species-specific qPCR assay first detected North American beavers in the dataset at 7,226 years ago (7.2 ka) in Jenny Lake and intermittently thereafter (10/14 samples) until present (fig 2). Beavers were first detected in Taggart Lake at 5,939 years ago (5.9 ka) and detected continuously from 5.2 ka until present. There was one detection gap in Taggart Lake at 5.5 ka. Mammalian sequencing with the 16SmammP007 barcode yielded sequences assigned to North American beavers in four samples - three in Jenny Lake and one in Taggart Lake - all of which also had positive qPCR assay beaver detections. Beavers were not detected in Lake Solitude, nor any of the negative controls with either the 16SmammP007 barcode or the qPCR assay.

**Figure 2:**
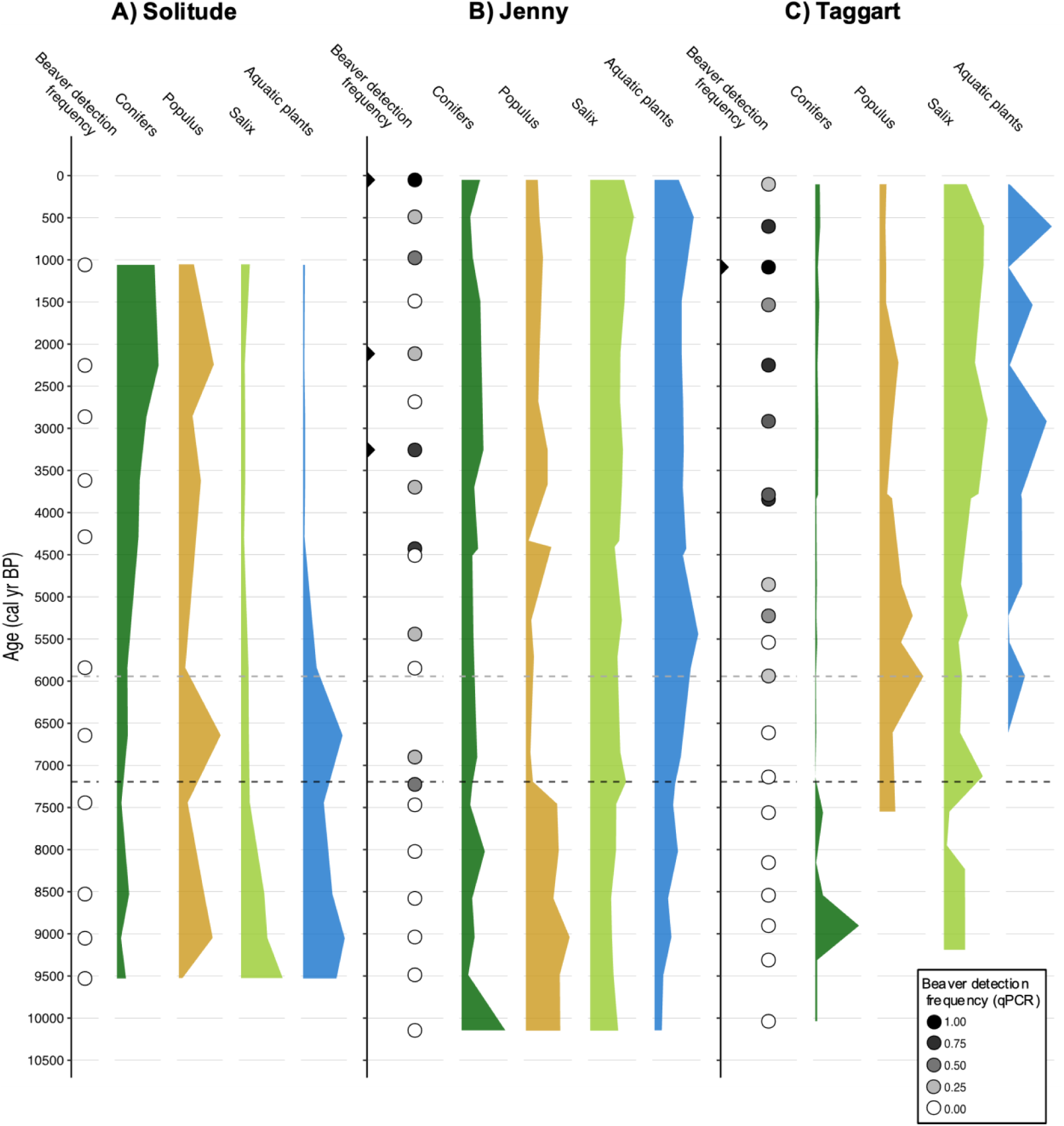
sedaDNA results from three Teton lakes: **a)** Lake Solitude, **b)** Jenny Lake, **c)** Taggart Lake. Leftmost panel for each subfigure indicates frequency of beaver detection across five replicates per sample from the species-specific qPCR assay with darker gray indicating higher detection frequency; black triangles indicate positive beaver detection with the 16SmammP007 barcode. From left to right remaining panels indicate relative frequency of reads per sample assigned to: conifers, *Populus*, *Salix*, and aquatic plants, scaled to maximum relative abundance per taxa per lake. Dashed horizontal lines indicate first appearance of beavers in Jenny Lake at 7.2 ka (black) and Taggart Lake at 5.9 ka (gray). Conifers includes all reads assigned to families *Cupressaceae* and *Pinaceae*; aquatic plants includes the genera: *Callitriche, Myriophyllum, Nuphar, Nymphaea,* and *Potamogeton*.

### 3.3. Vegetative trends

To evaluate the interaction between beavers and the local environment over time, we investigated trends in plant assemblages with a particular focus on taxa known to be associated with beavers (Fig. 2). In Taggart Lake, regional beaver arrival in the mid-Holocene is associated with a decrease in relative abundance of conifers and in increase in *Salix (*willows). The first detection of beavers in Taggart Lake coincides with the first detection of aquatic plants, which persist thereafter and become more abundant in the later Holocene. The first detection of *Populus* (e.g., poplar, aspen, cottonwood) in Taggart Lake slightly precedes the first regional detection of beavers; *Populus* relative abundance in Taggart Lake peaks at 5.9 ka when beavers are first locally detected and remains persistent throughout the remainder of the Holocene. *Salix* and aquatic plants also increase in Jenny Lake after beavers are first detected, similar to Taggart although to a lesser extent. Trends in conifers relative to beaver arrival in Jenny Lake are less clear and - in contrast to Taggart Lake - relative abundance of *Populus* is continuously high in the early Holocene, declining sharply at 7.2 ka when beavers are first detected, and increasing again in the later Holocene. Lake Solitude displays nearly opposite taxonomic trends to the lower elevation lakes, with initially high levels of aquatic plants, *Salix,* and *Populus* declining in the mid Holocene, coinciding with an increase in conifers.

We measured vegetation alpha diversity in each *trn*L sample using observed taxonomic richness and Shannon’s and Chao1 diversity indices. Shannon’s diversity index considers taxonomic evenness as well as richness, which effectively skews away from rarer taxa in the dataset; on the other hand, Chao1 is a nonparametric method that skews towards rare taxa (Kim et al. 2017). Diversity generally increased over time in Taggart and Jenny Lakes, while remaining stable in Lake Solitude (Fig. 3). We compared average diversity as measured by these three indices before and after the first regional detection of beavers at 7.2 ka. Diversity was significantly higher after 7.2 ka across all three indices in Taggart Lake, and in two of the three indices for Jenny Lake. Shannon diversity decreased slightly but non-significantly in Jenny Lake after 7.2 ka indicating a slight decline in taxonomic evenness. Diversity was generally low in Lake Solitude and did not change significantly before and after 7.2 ka for observed richness or Chao1 diversity, however there was a significant decrease in Shannon diversity.

**Figure 3:**
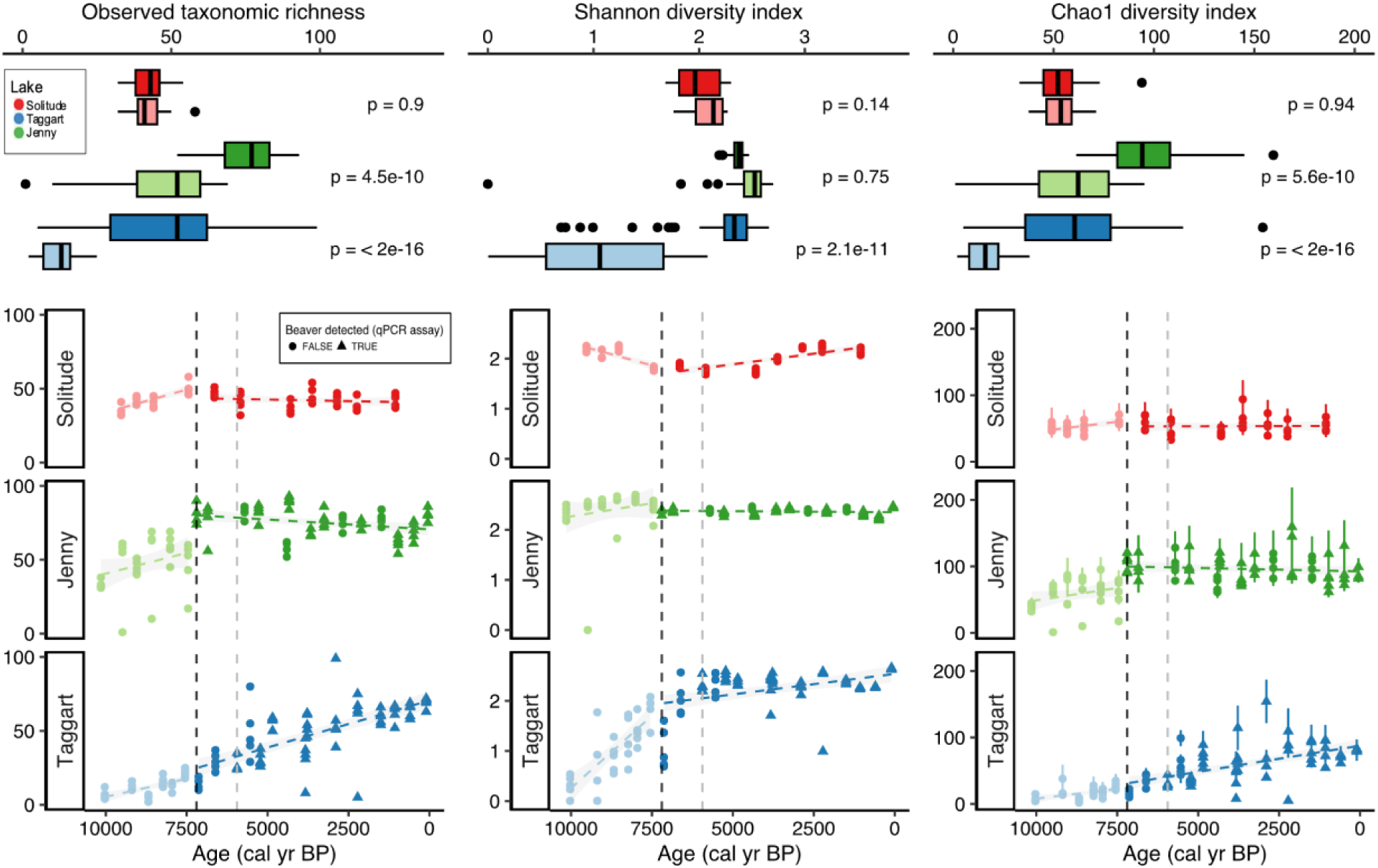
Alpha diversity indices for all trnL sample replicates over time (bottom) and means before and after beaver colonization at 7.2 ka compared (top) for each study lake (top to bottom: Solitude, Jenny, Taggart). Lighter shade indicates pre-beaver time period. Diversity indices left to right: Observed, Shannon, Chao1. Dashed vertical lines indicate first appearance of beavers in Jenny Lake at 7.2 ka (black) and Taggart Lake at 5.9 ka (gray).

Nonmetric multidimensional scaling (NMDS) analysis of the *trn*L relative abundance data yielded a minimum stress of 0.13, indicating good representation of the data by ordination. One outlier (the oldest sample from Taggart Lake) was removed from the NMDS plots for better visual representation. Samples plotting closer together in NMDS space indicates more similar plant communities. The biplot in Figure 4 demonstrates mid-Holocene regime shifts for all three lakes coinciding with the first detection of beavers; however, both Taggart and Jenny Lakes trend towards more positive MDS values, suggesting increasingly similar plant communities, whereas Lake Solitude shifts in the opposite direction. *Salix* and the majority of aquatic plant genera fall in the upper right quadrant of the NMDS plot with most of the post-beaver arrival Jenny and Taggart Lake samples, whereas most conifer genera plot on the right side of the plot with the Lake Solitude samples. When the MDS axes are plotted over time, Taggart Lake shows a strong shift in NMDS space associated with the first detection of beavers; changing from a negative temporal trend on MDS axis 1 before regional beaver arrival to a positive trend afterwards and more similar to Lake Solitude. MDS axis 2 shows a more consistent positive trend over time among all three lakes, although with a greater amplitude shift in Taggart Lake. PERMANOVA confirmed that plant communities differed significantly before and after the first beaver detection for all three lakes, considered both separately and together (Table 2).

**Figure 4:**
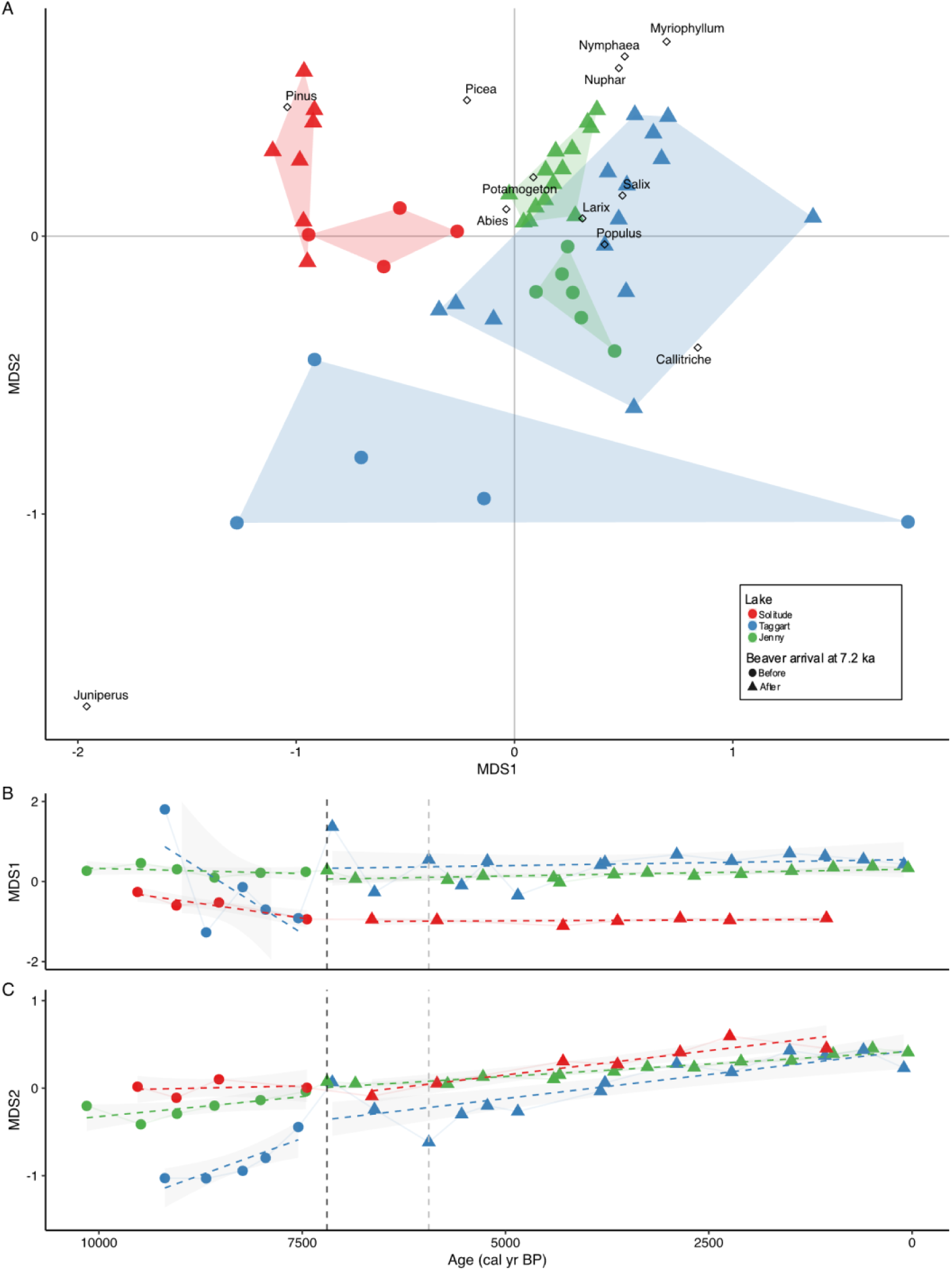
*trn*L beta diversity based on Jaccard similarity. **a)** Nonmetric multidimensional scaling (NMDS) axes 1 and 2 biplot with trnL samples (colored); and beaver-associated genera as in figure 2 (diamonds). **b)** MDS axis 1 and **c)** MDS axis 2 over time; trend lines for each lake and time period. Sample colors indicate lake and shape indicates time period (before or after beaver colonization); dashed vertical lines indicate first appearance of beavers in Jenny Lake at 7.2 ka (black) and Taggart Lake at 5.9 ka (gray).

**Table 2:** PERMANOVA results comparing plant community based on Jaccard similarity before and after first regional detection of beavers at 7.2 ka.

| Lake | Degrees of freedom | Sum of Squares | R<sup>2</sup> | Pseudo-F | P-value |
| --- | --- | --- | --- | --- | --- |
| Solitude | 1 | 0.47 | 0.39 | 5.64 | 0.008* |
| Jenny | 1 | 0.53 | 0.39 | 11.61 | 0.001* |
| Taggart | 1 | 0.96 | 0.20 | 4.57 | 0.001* |
| Combined | 2 | 2.99 | 0.29 | 9.79 | 0.001* |

## 4. Discussion

Using sedaDNA techniques we investigated beaver presence and vegetation diversity over the last 10 ka in three lakes in Grand Teton National Park, a region with a dynamic and well-described paleoecological history. We detected beavers in 21 lake sediment core samples up to 7.2 ka with a species-specific probe-based qPCR assay developed by Smith and Goldberg (2022). Although this assay was developed for modern eDNA applications, the high rate of detection indicates that it is a sensitive method for detecting the past presence of beavers in ancient sediments. Despite a shorter average amplicon size optimized for ancient DNA, the mammalian metabarcoding assay (16SmammP007) was much less effective; we recovered sequences assigned to beavers in only four samples, with a maximum detection age of 3.3 ka. We also detected beavers with the qPCR assay in all four of these samples, supporting its accuracy.

Using sedimentary analyses, Persico and Meyer (2013) found sporadic evidence of beaver activity in multiple stream beds in GTNP in the early Holocene, with more consistent detection in the later Holocene. These findings largely agree with our results here, lending support to the validity of this novel sedaDNA methodology. In particular, our first detection of beaver DNA at Taggart Lake at ∼5.9 ka is in close alignment with the ∼6 ka reported age of the oldest beaver-pond deposits identified along Beaver Creek, the lake’s outflow stream (Persico and Meyer, 2013). Additionally, many of the beaver detection gaps in the Jenny Lake record are temporally similar to those found by Persico and Meyer (2013), suggesting that these are real absences related to climatic and ecological changes. Furthermore, we tended to detect beavers at higher within-sample rates in time periods where Persico and Meyer also found highest levels of beaver activity.

Persico and Meyer (2013) reported two instances of beaver activity in the region at 10 ka and 8 ka, earlier than we detected beavers in this study. It’s possible that at this time, either beaver presence was too sparse and/or sporadic to be detected in our sampling, or that beavers were not active in the Taggart or Jenny Lake catchments despite being active in nearby streams. It is also possible that the qPCR assay was limited by DNA degradation in older samples. As such, the 7.2 ka beaver arrival time may reflect a methodological limit of detection rather than a biological reality. However, a shorter (60-84 bp) mammalian metabarcode did not identify beavers in any older samples, supporting the validity of the assay results. Future studies will explicitly test the temporal limits of this assay and potentially optimize shorter assays that may be more suitable for older and more degraded samples.

Our results suggest that during the Holocene, beavers first arrived to the Jenny Lake ecosystem no later than 7.2 ka and to Taggart Lake 5.9 ka, and were at least intermittently present in Jenny Lake throughout the remainder of the Holocene but were continuously present in Taggart Lake from 5.2 ka to present. The transition to mostly non-glacial conditions in the Tetons occurred by the start of the Holocene, approximately 11.5 ka, as evidenced by multiple lake sediment indicators (Larsen et al. 2016). The mid-Holocene was a time of environmental change in the Tetons, with a trend toward increased winter precipitation and cooler summers driving alpine glacier growth and raising regional moisture balance beginning around 6 ka (Larsen et al. 2020). Wetter conditions may have made the Cascade and Avalanche Canyons more hospitable to beavers, or increased riparian connectivity between the nearby Snake River and these lake systems, facilitating beaver movement into these watersheds. While beavers were historically present in high abundance throughout North America, the relatively late establishment of beavers in this region following deglaciation suggests that the spatial dynamics of beavers at the local scale may be quite complex.

Beavers appear to have arrived to the Taggart Lake watershed approximately 1.3 ka later than Jenny Lake, but were thereafter more persistent, with continuous detection in Taggart Lake from 5.2 ka to present while beavers were never continuously detected for more than ∼1.3 ka in Jenny Lake. Given the close proximity and similar geomorphology of these two lake systems, it is possible that these discrepancies represent a difference in DNA concentration and/or preservation rather than a true biological difference. Jenny Lake is much larger and deeper than Taggart Lake and has multiple sources of inflow; any eDNA in the system would therefore be more dilute and less detectable. Beavers are less likely to occupy lakes than rivers (Slough and Sadleir 1977), so it is likely that the DNA signal detected here was transported into the terminal lakes from their tributaries, further diluting the DNA signal. Cascade Canyon is longer and wider than Avalanche Canyon, providing more opportunities for DNA dilution. However, similar detection dynamics found by Persico and Meyer (2013) suggest that we may instead be documenting fine scale spatial and temporal dynamics of beaver activity in this region, with detection gaps in Jenny Lake corresponding closely to periods of reduced regional beaver activity identified by Persico and Meyer (2013) and attributed to drought.

Mid-Holocene plant community regime shifts are apparent in all three lakes coincident with beaver arrival. Based on relative abundance trends and beta diversity, Taggart Lake shows the strongest evidence of a mid-Holocene regime shift associated with beaver arrival, moving from a conifer-dominant to a more riparian system. Beaver-associated plants were either sporadically present (*Populus* and *Salix*) or absent (aquatic plants) until beaver arrival, and then consistently present thereafter. Alpha diversity also significantly increased after beaver arrival across all measures - consistent with predictions based on modern studies of how beavers influence plant diversity (eg. Naiman et al. 1999, Wright et al. 2002). The sustained detection of beavers in Taggart Lake from 5.2 ka until present suggests that beaver ecological engineering may have manipulated the environment in/around Taggart Lake enough to allow them to persist through periods of drought in the late Holocene that appear to have greatly reduced beaver abundance in nearby areas (Persico and Meyer 2009, 2013). The vegetation trends support this hypothesis, with beaver food sources *Populus*, *Salix,* and aquatic plants becoming much more consistent in Taggart Lake after beaver arrival - although aquatic plants undergo periods of decline at 2 and 1 ka, presumably as a result of these droughts. Beavers are known to “garden” preferred food species by altering local conditions in their favor (Naiman et al. 1988, Nummi 1989). The persistence of beavers and riparian plant communities through extended late Holocene droughts is an encouraging finding for beaver restoration, as it suggests that beaver activity may be able to maintain highly resilient watersheds that could provide refugia for plants and animals as the climate changes.

Jenny Lake shows largely similar vegetative trends as Taggart Lake but to a lesser degree. A notable difference is a sharp decrease in poplar relative abundance coinciding with beaver arrival. We can speculate that as a favored food source poplars were initially depleted by beaver arrival, but we do not have sufficient evidence to confirm this. We attribute the differences in plant community trends between Jenny and Taggart Lakes to the relative sizes of these two systems. Jenny Lake is much larger and deeper and captures a larger area, indicating a different set of controlling factors for both sedaDNA deposition and the aquatic and terrestrial communities. It is possible that as a smaller system, Taggart Lake is more sensitive to change and beavers therefore have a stronger effect on structuring the plant community. This may explain why beavers persist in Taggart Lake while disappearing from Jenny Lake during periods of presumed drought or other ecological stress.

Consistent with our predictions, we found no evidence of beavers in Lake Solitude, which is located near treeline in a much steeper and higher elevation cirque catchment. While beavers are capable of habitating high elevations and gradients, this environment represents more marginal habitat (McComb et al. 1990, Gurnell 1998). Despite no evidence of beavers, Lake Solitude also demonstrates a mid-Holocene vegetation regime shift albeit in an opposite direction from the lower elevation lakes, with increased conifer abundance and decreased riparian taxa. It is likely that this trend is attributable to some combination of decreased summer temperatures and increased in winter precipitation occurring at this time (e.g. Larsen et al., 2020). These changes would have created harsher conditions at high elevations while increasing the water available at lower elevations. Despite taxonomic compositional change, Lake Solitude showed little change in taxonomic richness over time, in contrast to the lower elevation lakes. This could be taken as evidence that beavers are driving these trends in richness, but it could also be that many taxa are limited by the altitude and generally harsh environment of Lake Solitude.

Taken together, the metabarcoding results of these three lakes suggests that a climatic shift in the mid-Holocene facilitated beaver establishment in the Jenny Lake and Taggart Lake catchments and likely contributed to coincident changes in the plant community. Paleoclimatic records indicate that regional winter precipitation and consequently lake levels increased at this time (Shuman and Serravezza 2017, Larsen et al. 2020). It is difficult to determine to what degree the mid-Holocene regime shifts apparent in these plant communities are attributable to beaver activity, rather than climatic shifts occurring at the time simply facilitated beaver establishment as well as plant community changes. We suggest that replicating this study at other locations in North America with similarly well-described paleoclimate histories will provide a clearer picture of the role of beavers in structuring local ecosystems throughout the Holocene.

We found that a species-specific qPCR assay applied to sedimentary samples is a powerful and reliable molecular method for detecting the past presence of beavers at the watershed scale in the absence of physical evidence. qPCR is faster, less expensive, and more analytically straightforward than other ancient eDNA methodologies such as metabarcoding or shotgun sequencing. The novel application of this molecular tool provides the opportunity to detect past beaver activity in a wide variety of settings without relying on sparsely distributed physical fossil or sedimentological evidence. A clearer picture of when and where this significant ecological engineer was active in the past can provide key insights as to how beavers may contribute to landscapes and ecosystem development. Furthermore, understanding the past temporal and spatial distribution of beavers can inform restoration and conservation efforts and help land managers better understand the effects of beaver engineering over long time scales and through changing climates.

## 5. Conclusions

Using a species-specific qPCR assay, we detected beaver sedaDNA in lake sediment samples up to 7.2 ka years old, demonstrating a sensitive method for documenting the historic presence of beavers in a watershed without physical evidence. Our findings show at least 5,000 years of continuous beaver presence in the Teton mountain range at Grand Teton National Park, suggesting that this important ecosystem engineer is an established and integral part of the local landscape. Our results largely agree with evidence of beaver activity previously established from sedimentary deposits in the region, supporting our conclusions and suggesting that sedaDNA is capable of reconstructing fine-scale spatial and temporal dynamics of past beaver activity. Our results suggest that beavers colonized the Taggart and Jenny Lake watersheds in the mid-Holocene. Evidence of regime shifts in the local plant community co-occurred with the establishment of beavers although questions remain as to what degree beavers were driving versus responding to local ecosystem dynamics. Although beavers appeared to be absent or greatly reduced in the Jenny Lake system during periods of inferred drought in the late Holocene, they remained consistently present in the Taggart Lake catchment, perhaps as a result of intensive ecological engineering at the local scale. The continuous persistence of beavers at this site indicates that under certain conditions, beavers may be able to maintain wetlands through periods of climatic stress, providing important refugia and buffering the effects of climate change at the local scale. A better understanding of regional beaver dynamics during periods of historic climate change will provide a clearer picture of how common this may be and what conditions beavers need in order to maintain continuous presence. These results shed light on the role of beavers in North American paleoecosystems and may help managers more effectively deploy beaver engineering as a climate mitigation strategy.

## Acknowledgements

We thank Simeon Caskey and other NPS staff for facilitating this research and for logistical support. We thank A. Blumm, J. Raberg, S. Pendleton, D. Weber, N. Reitman, X. Wang, J. Von Eggers, H. Bender, and S. Sacco for field and lab assistance. Initial core processing and analyses were performed at the NSF-shared LacCore facility, University of Minnesota.

