## Supplemental Information for "Ancient sedimentary DNA shows more than 5000 years of continuous beaver occupancy in Grand Teton National Park"

### Supplementary Information

#### 1. Sediment core methods and geochronology

Sediment cores were collected from Taggart Lake and Jenny Lake in 2019 and 2021, respectively, using two types of coring systems deployed on cables from the lake surfaces (Table S1). At both lakes, long cores were collected through repeated 2-m drives using a Uwitec coring system and a surface core was collected using a modified Nesje corer. Sediment cores were collected from Lake Solitude in 2020 through repeated 1-m drives using a lance-driven Livingstone-type corer and a Bolivia-type surface corer. All cores from the three lakes were collected from the stable lake-ice surface, packaged in the field, and transported to the Sedimentology and Paleoclimate lab at Occidental College for initial core processing and description. Core sections were split lengthwise into working and archive halves and core halves photographed using a linescan core imager prior to storage in a climate-controlled core repository.

Age control of Teton lake sediments was established using Accelerated Mass Spectrometry (AMS 14C) radiocarbon dating and tephrochronology (Fig. S1; Table S2). In all cases, samples of terrestrial plant material (wood, charcoal, conifer needles) selected for AMS 14C were pretreated and measured at the W.M. Keck Carbon Cycle AMS Laboratory, University of California, Irvine. One prominent rhyolitic tephra layer was visually observed in core sections from all three lakes and assigned to the Mazama tephra bed based on analyses described by Larsen et al. (2016). The compiled list of all radiocarbon and tephra ages from all lake cores is presented in Table S2 below. The age models for all three lake sediment sequences are presented in Figure S1 below. Note, the age model for core JEN21-1 from Jenny Lake was previously established by Larsen et al. (2024).

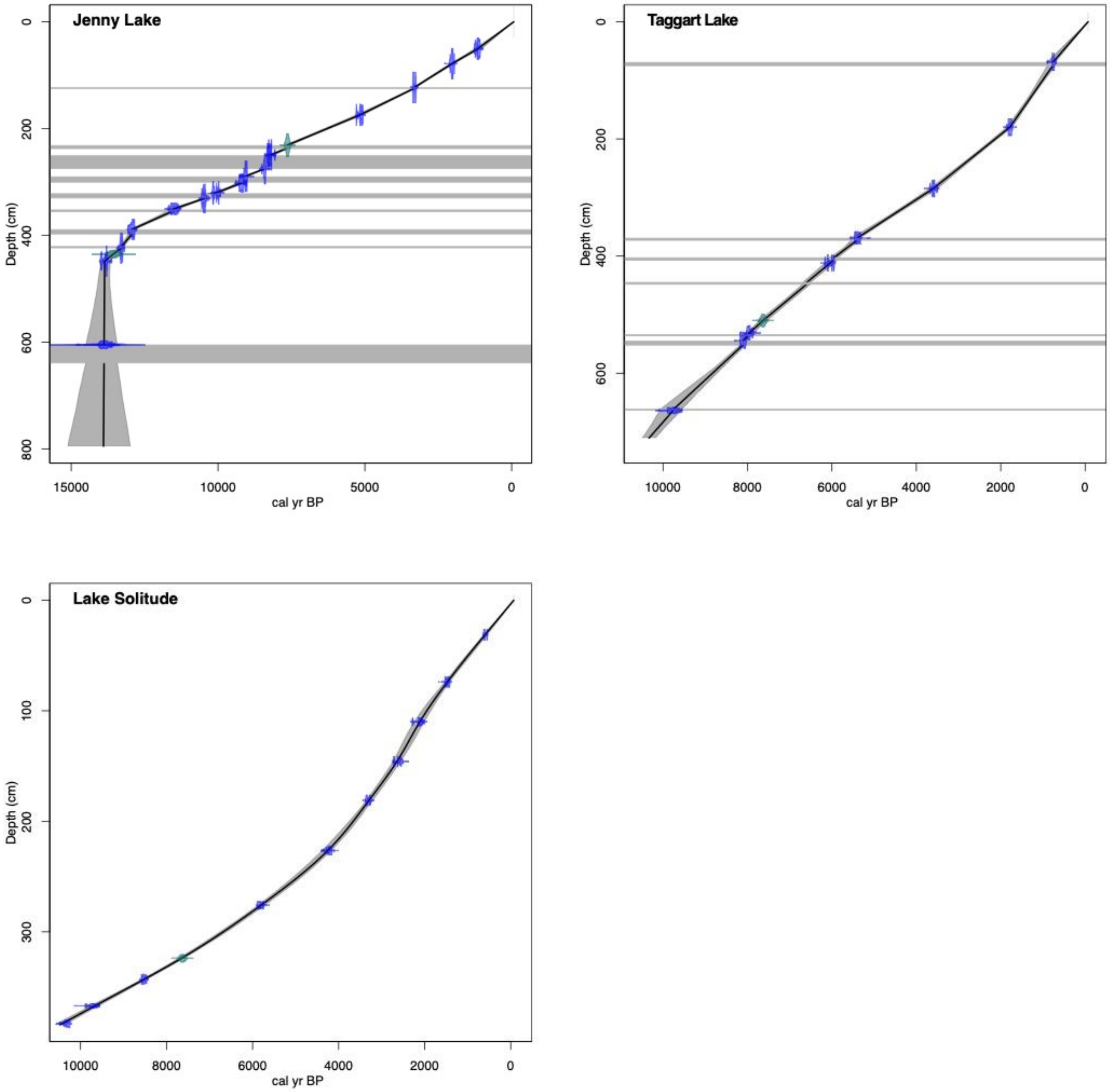


**Supplementary Figure 1: Age models for Teton lake cores recovered from Jenny Lake, Taggart Lake, and Lake Solitude.** Core age models were generated with a linear interpolation (Jenny Lake, Taggart Lake) and smooth spline interpolation (Lake Solitude) of AMS radiocarbon and tephra control points using CLAM code for R software. Horizontal grey shade bars in Jenny Lake and Taggart Lake age models represent turbidite deposits.

**Supplementary Table 1: Teton lake sediment core metadata.**

| Core ID | Latitude | Longitude | Water depth (m) | Core length (m blf) |
|---|---|---|---|---|
| JEN21-1 | 43.7635° | -110.7308° | 68.0 | 7.9 |
| TAG19-1A | 43.7053° | -110.7561° | 9.3 | 12 |
| SOL21-1 | 43.7927° | -110.8443° | 10.4 | 6.2 |

**Supplementary Table 2: Sediment core radiocarbon and tephra ages with calibrated 2 sigma error ranges.** (Note: samples marked with * were considered outliers and excluded from age models.)

| Sample Lab ID | Core ID | Cumulative depth (cm) | Material dated | Uncalibrated age ($^{14}C$ yr BP) | Error (yr) | Calibrated age (cal yr BP) (2sigma) |
|---|---|---|---|---|---|---|
| UCIAMS257786 | JEN21-1 | 50.5 | needle | 1225 | 40 | 1146 (1059—1275) |
| OS162108 | JEN21-1 | 78.5 | needle | 2070 | 25 | 2031 (1944—2116) |
| CURL23045 | JEN21-1 | 123 | wood fragments | 3105 | 20 | 3328 (3246—3379) |
| CURL23053 | JEN21-1 | 174.5 | needle | 4520 | 20 | 5152 (5051—5306) |
| Mazama | JEN21-1 | 231 | tephra | | | 7631 (7579—7683) |
| UCIAMS269931 | JEN21-1 | 249.5 | charcoal | 7385 | 25 | 8204 (8041—8324) |
| CURL24880 | JEN21-1 | 250 | needle | 7445 | 25 | 8264 (8189—8339) |
| UCIAMS269933 | JEN21-1 | 275.5 | plant material | 7620 | 40 | 8409 (8361—8519) |
| OS162109 | JEN21-1 | 289.5 | plant material | 8120 | 30 | 9061 (8993—9192) |
| UCIAMS164819 | JEN21-1 | 302 | needle | 8240 | 25 | 9208 (9033—9399) |
| UCIAMS164820 | JEN21-1 | 320.5 | needle | 8940 | 30 | 10058 (9911—10201) |
| UCIAMS172641 | JEN21-1 | 331 | needle | 9285 | 25 | 10478 (10308—10574) |
| CURL23041 | JEN21-1 | 350.5 | plant material | 10010 | 30 | 11492 (11314—11698) |
| CURL23062 | JEN21-1 | 389 | wood fragments | 11000 | 35 | 12916 (12828—13070) |
| UCIAMS287129 | JEN21-1 | 424 | wood fragments | 11430 | 30 | 13301 (13183—13410) |
| Glacier Peak | JEN21-1 | 435 | tephra | | | 13560 (13410—13710) |
| UCIAMS141222 | JEN21-1 | 448.5 | plant material | 11945 | 35 | 13835 (13612—14021) |
| CURL24945 | JEN21-1 | 605 | charcoal | 11840 | 410 | 13839 (12928—15081) |
| CURL24933 | JEN21-1 | 605 | needle | 11960 | 120 | 13845 (13520—14081) |
| CURL28109 | TAG19-1A | 68.5 | needle and wood fragments | 860 | 25 | 757 (689—897) |
| CURL28105 | TAG19-1A | 180 | seed | 1880 | 25 | 1784 (1718—1865) |
| CURL28101 | TAG19-1A | 284.5 | needle | 3360 | 25 | 3594 (3491—3687) |
| OS160703 | TAG19-1A | 369 | plant material | 4670 | 35 | 5398 (5315—5474) |
| CURL28106 | TAG19-1A | 412 | wood fragment | 5260 | 25 | 6048 (5934—6177) |
| Mazama | TAG19-1A | 510 | tephra | | | 7631 (7579—7683) |
| OS160704 | TAG19-1A | 531 | needle and wood fragments | 7140 | 60 | 7960 (7797—8160) |
| UCIAMS287131 | TAG19-1A | 544.5 | wood | 7325 | 20 | 8100 (8035—8179) |
| CURL28108 | TAG19-1A | 552.5 | wood | 7660 | 30 | *8440 (8389—8536) |
| OS159524 | TAG19-1A | 663.5 | plant material | 8760 | 45 | 9752 (9551—10110) |
| UCIAMS269930 | SOL20-1 | 31.25 | needle | 585 | 20 | 606 (542—642) |
| UCIAMS257814 | SOL20-1 | 47 | plant material | 4495 | 15 | *5166 (5048—5288) |
| OS159520 | SOL20-1 | 56.5 | wood | 1520 | 15 | *1386 (1355—1407) |
| CURL28094 | SOL20-1 | 74 | plant material | 1610 | 25 | 1476 (1412—1534) |
| OS159521 | SOL20-1 | 108 | seed | 2440 | 20 | *2474 (2360—2696) |
| OS160702 | SOL20-1 | 110 | needle fragments | 2140 | 30 | 2116 (2000—2299) |
| CURL28099 | SOL20-1 | 146 | needle | 2530 | 25 | 2617 (2496—2739) |
| CURL28118 | SOL20-1 | 181 | needle and wood fragments | 3105 | 25 | 3322 (3241—3381) |
| CURL28113 | SOL20-1 | 226.5 | needle fragments | 3830 | 25 | 4222 (4099—4396) |
| CURL28100 | SOL20-1 | 276 | needle | 5055 | 25 | 5824 (5737—5899) |
| Mazama | SOL20-1 | 324 | tephra | | | 7631 (7579—7683) |
| UCIAMS252974 | SOL20-1 | 343 | needle fragments and wood | 7750 | 20 | 8527 (8453—8591) |
| CURL28116 | SOL20-1 | 367 | needle fragments | 8745 | 35 | 9717 (9555—9892) |
| CURL28107 | SOL20-1 | 383 | needle fragments | 9165 | 35 | 10319 (10238—10485) |

**Supplementary Table 3: sedaDNA subsample information.**

| Extraction ID | Lake | Core ID | Section depth (cm) | Cumulative depth (cm) | Calibrated age (cal yr BP) |
|---|---|---|---|---|---|
| DNB031.11 | Jenny | JEN21-1A-1N-1 | 2 | 5 | 50 |
| DNB031.10 | Jenny | JEN21-1A-1N-1 | 18 | 21 | 437 |
| SC126 | Jenny | JEN21-1B-1U-1 | 22 | 23 | 486 |
| DNB031.09 | Jenny | JEN21-1A-1N-1 | 38 | 41 | 922 |
| SC125 | Jenny | JEN21-1B-1U-1 | 42 | 43 | 970 |
| DNB031.08 | Jenny | JEN21-1A-1N-1 | 58 | 61 | 1482 |
| SC124 | Jenny | JEN21-1B-1U-1 | 62 | 63 | 1545 |
| DNB031.07 | Jenny | JEN21-1A-1N-1 | 78 | 81 | 2104 |
| SC123 | Jenny | JEN21-1B-1U-1 | 82 | 83 | 2161 |
| DNB031.06 | Jenny | JEN21-1A-1N-1 | 98 | 101 | 2680 |
| SC122 | Jenny | JEN21-1B-1U-1 | 102 | 103 | 2737 |
| DNB031.05 | Jenny | JEN21-1A-1N-1 | 118 | 121 | 3256 |
| SC121 | Jenny | JEN21-1B-1U-1 | 122 | 123 | 3313 |
| SC120 | Jenny | JEN21-1B-1U-2 | 5 | 134 | 3665.5 |
| DNB033.05 | Jenny | JEN21-1A-1N-2 | 5 | 152 | 4333.5 |
| SC119 | Jenny | JEN21-1B-1U-2 | 25 | 154 | 4407.5 |
| DNB033.04 | Jenny | JEN21-1A-1N-2 | 30 | 177 | 5276.5 |
| DNB033.03 | Jenny | JEN21-1A-1N-2 | 40 | 187 | 5712.5 |
| DNB033.02 | Jenny | JEN21-1B-2U-01 | 1 | 213 | 6846 |
| DNB033.01 | Jenny | JEN21-1B-2U-01 | 9 | 221 | 7195 |
| SC118 | Jenny | JEN21-1B-2U-01 | 15 | 227 | 7456 |
| SC117 | Jenny | JEN21-1B-2U-01 | 33 | 245 | 8012 |
| SC116 | Jenny | JEN21-1B-2U-01 | 67 | 279 | 8579 |
| SC137 | Jenny | JEN21-1B-2U-01 | 77 | 289 | 9040 |
| SC136 | Jenny | JEN21-1B-2U-01 | 96 | 308 | 9484 |
| SC104 | Jenny | JEN21-1B-2U-01 | 104 | 316 | 9859 |
| SC115 | Jenny | JEN21-1B-2U-01 | 109 | 321 | 10150 |
| SC083 | Solitude | SOL20-1B-1L-1 | 3 | 53 | 1053 |
| SC084 | Solitude | SOL20-1B-1L-1 | 69 | 119 | 2242 |
| SC085 | Solitude | SOL20-1B-2L-1 | 11 | 158 | 2853 |
| SC086 | Solitude | SOL20-1B-2L-1 | 51 | 198 | 3622 |
| SC087 | Solitude | SOL20-1B-2L-1 | 81 | 228 | 4292 |
| SC088 | Solitude | SOL20-1B-3L-1 | 27 | 277 | 5842 |
| SC089 | Solitude | SOL20-1B-3L-1 | 49 | 299 | 6643 |
| SC090 | Solitude | SOL20-1B-3L-1 | 69 | 319 | 7442 |
| SC091 | Solitude | SOL20-1B-3L-1 | 93 | 343 | 8524 |
| SC092 | Solitude | SOL20-1B-4L-1 | 7 | 354 | 9047 |

| | | | | | |
|---|---|---|---|---|---|
| SC093 | Solitude | SOL20-1B-4L-1 | 17 | 364 | 9525 |
| DNB031.04 | Taggart | TAG19-2A-1L-1 | 14 | 14 | 101 |
| DNB031.03 | Taggart | TAG19-2A-1L-1 | 20 | 20 | 173 |
| DNB030.11 | Taggart | TAG19-1A-1U-1 | 6 | 21 | 185 |
| DNB031.02 | Taggart | TAG19-2A-1L-1 | 30 | 30 | 295 |
| DNB031.01 | Taggart | TAG19-2A-1L-1 | 45 | 45 | 476 |
| DNB032-11 | Taggart | TAG19-1A-1U-1 | 40 | 55 | 598 |
| DNB032-10 | Taggart | TAG19-1A-1U-1 | 66 | 81 | 829 |
| DNB030.10 | Taggart | TAG19-1A-1U-1 | 91 | 106 | 1070.5 |
| DNB032-09 | Taggart | TAG19-1A-1U-1 | 136 | 151 | 1505.5 |
| DNB030.09 | Taggart | TAG19-1A-1U-2 | 40 | 205 | 2218.5 |
| DNB032-08 | Taggart | TAG19-1A-2U-1 | 5 | 244 | 2894 |
| DNB032-07 | Taggart | TAG19-1A-2U-1 | 54 | 293 | 3777 |
| DNB030.07.08 | Taggart | TAG19-1A-2U-1 | 56.5 | 295.5 | 3831 |
| DNB032-06 | Taggart | TAG19-1A-2U-1 | 104 | 343 | 4848 |
| DNB030.06 | Taggart | TAG19-1A-2U-1 | 121.5 | 360.5 | 5223 |
| DNB032-05 | Taggart | TAG19-1A-2U-1 | 141 | 380 | 5539 |
| DNB030.05 | Taggart | TAG19-1A-2U-2 | 19.5 | 402.5 | 5943 |
| DNB032-04 | Taggart | TAG19-1A-3U-1 | 5 | 449 | 6611 |
| DNB030.04 | Taggart | TAG19-1A-3U-1 | 36 | 480 | 7130 |
| DNB032-03 | Taggart | TAG19-1A-3U-1 | 61 | 505 | 7548 |
| DNB030.03 | Taggart | TAG19-1A-3U-1 | 87 | 531 | 7950 |
| DNB032-02 | Taggart | TAG19-1A-3U-1 | 116 | 560 | 8232 |
| DNB030.02 | Taggart | TAG19-1A-3U-1 | 146 | 590 | 8681 |
| DNB032-01 | Taggart | TAG19-1A-3U-2 | 31 | 624 | 9189 |
| DNB030.01 | Taggart | TAG19-1A-4U-1 | 42 | 686 | 10030.5 |
